# Increased Susceptibility of WHIM Mice to Papillomavirus-induced Disease is Dependent upon Immune Cell Dysfunction

**DOI:** 10.1101/2023.11.15.567210

**Authors:** Wei Wang, Ali Pope, Ella Ward-Shaw, Darya Buehler, Francoise Bachelerie, Paul F. Lambert

## Abstract

Warts, Hypogammaglobulinemia, Infections, and Myelokathexis (WHIM) syndrome is a rare primary immunodeficiency disease in humans caused by a gain of function in CXCR4, mostly due to inherited heterozygous mutations in *CXCR4*. One major clinical symptom of WHIM patients is their high susceptibility to human papillomavirus (HPV) induced disease, such as warts. Persistent high risk HPV infections cause 5% of all human cancers, including cervical, anogenital, head and neck and some skin cancers. WHIM mice bearing the same mutation identified in WHIM patients were created to study the underlying causes for the symptoms manifest in patients suffering from the WHIM syndrome. Using murine papillomavirus (MmuPV1) as an infection model in mice for HPV-induced disease, we demonstrate that WHIM mice are more susceptible to MmuPV1-induced warts (papillomas) compared to wild type mice. Namely, the incidence of papillomas is higher in WHIM mice compared to wild type mice when mice are exposed to low doses of MmuPV1. MmuVP1 infection facilitated both myeloid and lymphoid cell mobilization in the blood of wild type mice but not in WHIM mice. Higher incidence and larger size of papillomas in WHIM mice correlated with lower abundance of infiltrating T cells within the papillomas. Finally, we demonstrate that transplantation of bone marrow from wild type mice into WHIM mice normalized the incidence and size of papillomas, consistent with the WHIM mutation in hematopoietic cells contributing to higher susceptibility of WHIM mice to MmuPV1-induced disease. Our results provide evidence that MmuPV1 infection in WHIM mice is a powerful preclinical infectious model to investigate treatment options for alleviating papillomavirus infections in WHIM syndrome.

**AUTHOR SUMMARY:** Mice carrying the same gain-of-function mutation in the gene *CXCR4* that is present in human patients suffering from the Warts, Hypogammaglobulinemia, Infections, and Myelokathexis (WHIM) syndrome were previously created to understand the biology underlying this syndrome and to develop better means for treating WHIM patients. WHIM mice display neutropenia and lymphopenia symptoms as do WHIM patients. One of the key features of the WHIM syndrome in humans is increased susceptibility to infections by human papillomaviruses (HPV) with the majority of WHIM patients experiencing persistent warts and some developing anogenital cancers, both caused by HPVs. In this study we use a mouse papillomavirus, MmuPV1, which is a model for HPV infection in humans, to ask if the WHIM mice are more susceptible to infection and to understand why. We demonstrate that WHIM mice are more susceptible to MmuPV1-induced disease and that correcting the neutropenia and lymphopenia by bone marrow transplantation was effective at decreasing susceptibility to MmuVP1 induced disease. Our data support WHIM mice as a disease model for WHIM syndrome for future investigations on curative treatment options.

## INTRODUCTION

WHIM syndrome is a rare immunodeficiency disease characterized by <u>W</u>arts, <u>H</u>ypogammaglobulinemia, Infections and <u>M</u>yelokathexis. WHIM patients are characterized by neutropenia, lymphopenia and high susceptibility to repeated bacterial infections, and HPV pathogenesis manifesting as skin, mucosal, oral, and genital warts or intraepithelial neoplasia that may progress to cancer (1). Past research identified dominant mutations in the *CXCR4* gene that are inheritable and result in a gain-of-function of CXCR4 protein function, which causes the symptoms of the WHIM syndrome (2). These mutations result in an early truncation of CXCR4 cytoplasmic tail, leading to impaired CXCR4 desensitization including the decreased internalization of cell surface CXCR4 upon CXCL12 binding (3). CXCL12, the only known chemokine ligand for CXCR4, is abundantly expressed by bone marrow stromal cells, such that this signaling axis regulates hematopoietic stem cells (HSCs) homing from blood to bone marrow niches (4). CXCR4 is also responsible for maintaining HSC quiescence and controlling their proliferation (5). Most leukocytes and myeloid cells also express surface CXCR4 (6). Consequently, the gain of function mutations in the CXCR4/CXCL12 signaling axis of WHIM syndrome patients, are associated with panleukopenia and neutrophil retention in bone marrow (7). Currently there are two CXCR4 antagonists in clinical trials for treating WHIM syndrome. Mavorixafor (X4P-001) is a selective small molecule CXCR4 inhibitor that is orally administered daily to WHIM patients. Upon long term treatment, one patient experienced 79% reduction in warts (8). Another promising CXCR4 antagonist for WHIM syndrome is Plerixafor (Mozobil, AMD3100). Plerixafor is an FDA-approved drug for mobilizing HSC to peripheral blood as a preparation for autologous transplant patients with non-Hodgkin’s lymphoma and multiple myeloma (9, 10). Plerixafor was shown to reduce epithelial hyperplasia in HPV transgenic mice (11) and correct leukopenia in WHIM patients (12, 13). In terms of a cure for WHIM syndrome, HSC transplantation remains the only option for now. However, there are mixed results reported for WHIM patients who received bone marrow transplant (14). Two out of seven studies reported that patients who underwent bone marrow transplant experienced severe disease complications, with one of them dying. The rest of the patients survived with full donor chimerism at long-term follow-up (median 6.7 years) (14). Due to these complications, bone marrow transplant has not been adopted as a standard for treating patients with WHIM syndrome but rather is used on a case-by-case basis.

Around 40-78% of WHIM patients develop widespread HPV-induced warts by the age of 30 and WHIM patients also have an increased likelihood of developing HPV-related malignancies compared to the general population (1, 15–18). It remains unclear whether prophylactic HPV vaccines will be effective at protecting WHIM patients against high-risk alpha HPVs associated with anogenital cancers and head and neck cancers targeted by these vaccines. In one study, vaccination of a WHIM patient was found to induce antibodies against the HPVs targeted; however, the titers of those antibodies were greatly reduced compared to that seen in the healthy population (18). This may reflect the fact that WHIM patients suffer from hypogammaglobulinemia. In addition, these HPV vaccines do not prevent infections by cutaneous HPVs that cause disfiguring warts, a common ailment for WHIM patients (19, 20). Therefore, further study is necessary to learn how to best prevent/treat HPV-associated disease in WHIM patients.

Papillomaviruses are species specific viruses. Therefore, studying HPV infection-induced pathogenesis in laboratory animals is not possible. In 2011, a papillomavirus that infects laboratory mice (Mus musculus) was described (21). Since then, this mouse papillomavirus, MmuPV1, has been found to infect both cutaneous and mucosal epithelia of immunocompetent FVB/N mice and to cause cancers in these same tissues in which HPVs cause cancer in humans (22–26). The ability of MmuPV1 to cause disease in mice is dependent upon its ability to evade immune detection (24, 25), and this mechanism of immune evasion appears to be shared with HPVs (27–29) and to contribute to resistance to immune checkpoint blockade immunotherapy in humans (30). MmuPV1, therefore, provides us a powerful new infection-based preclinical model for studying HPV biology.

A knock-in mouse strain with a WHIM syndrome-related mutation in *Cxcr4* gene (CXCR4^1013^) was created to study underlying mechanisms of *CXCR4* mutation caused WHIM pathogenesis (31). Like WHIM patients, mice bearing the *CXCR4* mutation display neutropenia and lymphopenia, while possessing normal bone marrow architecture and granulocyte lineage maturation (31). Using WHIM mice as a model, Murphy et al, recently suggested that WHIM allele inactivation may be a preferred strategy for genetic cure of WHIM syndrome (32). In this study, we asked if mice carrying the WHIM mutation (CXCR4^1013^) are increased in their susceptibility to disease induced by the mouse papillomavirus, MmuPV1, and whether we could correct for this increased susceptibility by bone marrow transplantation. We found this strain to indeed display increased susceptibility to MmuPV1-induced papillomatosis at a low dose of infection compared to wild type mice, and to display continued neutropenia and lymphopenia. We found that reconstituting wild type mice with bone marrow cells from WHIM mice conferred increased susceptibility to MmuPV1-induced warts, and, conversely, reconstituting WHIM mice with bone marrow cells from wild type mice normalized papilloma incidence to that seen in wild type mice. Our findings indicate that this mouse papillomavirus infection model for WHIM syndrome recapitulates the high susceptibility to HPV-induced warts in human WHIM patients, and that this increased susceptibility is conferred, at least in part, by alterations in the properties of immune cells carrying the CXCR4^1013^ mutation. This preclinical model should provide a useful platform for testing treatment options to prevent and/or cure papillomavirus infections in WHIM syndrome patients.

## RESULTS

### WHIM mice have increased susceptibility to MmuPV1-induced papillomatosis

We infected the skin on the ears of wild type (WT) mice (WT/WT), mice heterozygous for the WHIM mutant (M) allele (WT/M) and mice homozygous for the WHIM mutant allele (M/M), with three doses (viral genome equivalents: VGE) of MmuPV1 (10^8^ VGE, 10^7^ VGE and 10^6^ VGE). The incidence of MmuPV1-induced papillomatosis on FVB/N ear skin usually plateaus at 4 weeks post infection (24). Because skin MmuPV1 infection requires scarification to expose the basal layer of epithelium to virus (33), and because wound healing process takes approximately 3 weeks to resolve, the presence of ear papillomas was scored and size of papillomas was measured at 4 weeks post-infection. The incidence and size of papillomas (warts) were dependent upon MmuPV1 dose (Figure 1). At a high dose of infection (2x10^8^ VGE), the majority of infection sites on WT animals and all infection sites of WHIM mice (WT/M and M/M) developed warts by 4 weeks after infection (Figure 1A). The size of warts arising in WHIM mice (WT/M and M/M) were significantly larger than warts arising in WT mice (Figure 1B). At a low dose of MmuPV1 infection (10^6^ VGE), only 10% of infected sites in WT animals developed warts (Figure 1A). In contrast, over 50% of infected sites in mice heterozygous or homozygous for WHIM mutation developed warts (Figure 1A), with bigger sizes of warts compared to warts arising on WT mice (Figure 1B and 1C). Neither the incidence nor wart size were significantly different between heterozygous and homozygous WHIM mice (Figure 1), consistent with this mutant phenotype being dominant over wild type in mice, with no clear dosage effect from the CXCR4 mutation being observed. *In situ* hybridization of MmuPV1 DNA showed a significantly higher number of keratinocytes positive for MmuPV1 DNA in warts arising in WHIM mice compared to that in WT mice (Figure 1D), but there was not a significant difference in levels of expression of MmuPV1 RNA expression or viral proteins (E4, L1) (Figures 1E and 1F). This difference in MmuPV1 DNA level indicates that MmuVP1-infected WHIM keratinocytes have an advantage in supporting viral DNA replication.

**Figure 1.**
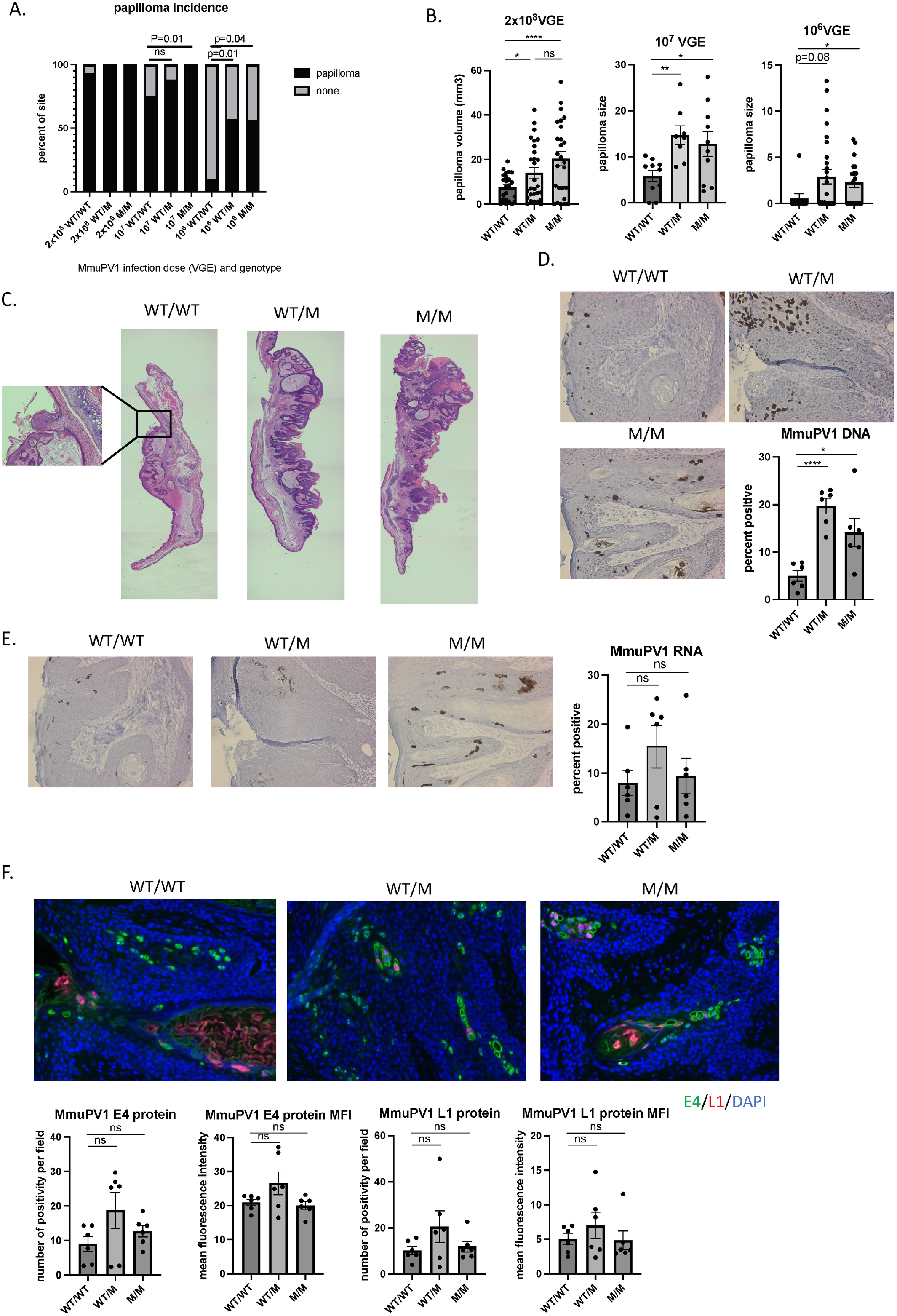
WHIM mice have increased susceptibility to MmuPV1-mediated papillomatosis. Both ears of wildtype (WT/WT) littermates, WHIM heterozygous mice (WT/M) and WHIM homozygous mice (M/M) were infected with 2x10^8^ VGE, 1X10^7^ VGE and 1x10^6^ VGE respectively. A) Incidence of papillomatosis was scored at 4 weeks post infection. Fisher’s exact test was used to compare between genotypes with the same dosage of infection. B) Wart size were measure at 4 weeks post infection. Student t test was used to compare wart size between genotypes. C) Representative 2.5x H&E images of warts at 4 weeks post infection from wildtype mice (WT/WT), WHIM heterozygous mice (WT/M) and WHIM homozygous mice (M/M). Four images from each wart were taken under 2.5x magnification and stitched together to cover the whole length of tissue. 10x image of the WT wart was shown next to it to show hyperplasia. D) Representative 20x images of MmuPV1 DNA in situ hybridization at 4 weeks post infection from wildtype mice (WT/WT), WHIM heterozygous mice (WT/M) and WHIM homozygous mice (M/M). Number of DNA positive cells and total nucleated cells were counted. Three 20x fields were averaged for one wart. Six warts from each group were quantified. Student t test was used. E) Representative 20x images of MmuPV1 RNA in situ hybridization at 4 weeks post infection from wildtype mice (WT/WT), WHIM heterozygous mice (WT/M) and WHIM homozygous mice (M/M). Number of DNA positive cells and total nucleated cells were counted. Three 20x fields were averaged for one wart. Six warts from each group were quantified. Student t test was used. F) Representative 20x images from MmuPV1 E4 and L1 protein immunofluorescence staining at 4 weeks post infection from wildtype mice (WT/WT), WHIM heterozygous mice (WT/M) and WHIM homozygous mice (M/M). Number of E4 or L1 positive signals were counted per field. Mean fluorescence intensity per field is quantified by ImageJ. Three 20x fields were averaged for one wart. Six warts from each group were quantified. Ns: not significant. *p<0.05, **p<0.01, ****p<0.001.

### WHIM mice have disturbed T cell distribution in the blood, skin-warts and bone marrow upon MmuPV1 infection

Blood was collected from animals with all three genotypes prior to MmuPV1 infection and 4 weeks post infection (2x10^8^ VGE, this dose was chosen because nearly all mice infected with this dose of virus show evidence of disease both in WT and WHIM mice at 4 weeks post-infection – see Figure 1A). Whole blood was subjected to complete blood count. As previously reported (31), at a steady state (pre-infection), WHIM mice carrying one or two CXCR4 mutations suffer from panleukopenia and had significantly lower counts of white blood cell counts (WBC), neutrophils and lymphocytes in their blood (Figure 2A). Following MmuPV1 infection, circulating levels of WBC, neutrophils and lymphocytes increased in WT animals (Figure 2A). These increases were modest in heterozygous (WT/M) and homozygous (M/M) mice following MmuPV1 infection (Figure 2A). We collected warts after 4 weeks post infection to quantify papilloma-infiltrating immune cells by flow cytometry. Warts arising in WHIM mice (WT/M and M/M) had fewer CD45+, CD4+ and CD8+ infiltrating immune cells, respectively, compared to warts arising in WT mice (Figure 2B). The reductions in subsets of infiltrating myeloid cells and B cells infiltrating the warts in WHIM mice were not significant (Figure 2B). The difference in the frequency of wart-infiltrating CD8+ cells in WHIM mice versus WT mice was confirmed by immunofluorescence staining of CD8+ and K14+ cells and by counting the number of intra-epithelial CD8+ cells in warts (white arrows, Figure 2C). In parallel, we looked at levels of immune cells in the bone marrow, because it has been documented that in WHIM mice there is an increased homing of certain immune cells to the bone marrow as a consequence of the gain-of-function mutation in CXCR4 (31). In naïve (uninfected) mice we observed significant increased abundance of neutrophils, CD4+ T cells and CD8+ T cells and a decreased abundance of B cells in the bone marrow of WHIM mice compared to WT mice (Figure 2D, top graphs). These differences were blunted in infected mice (Figure 2D, bottom graphs). These data demonstrate that WHIM mice experience prolonged neutropenia and lymphopenia even with papillomavirus infection. The observation that there is a highly significant (p<0.01) decrease specifically in CD8+ T cell infiltration within the papillomas arising in WHIM mice compared to in WT mice, but a much lesser difference for other immune cell types, raises the possibility that this deficit, in particular, is contributing to the higher incidence of papillomatosis and faster wart growth in WHIM mice.

**Figure 2.**
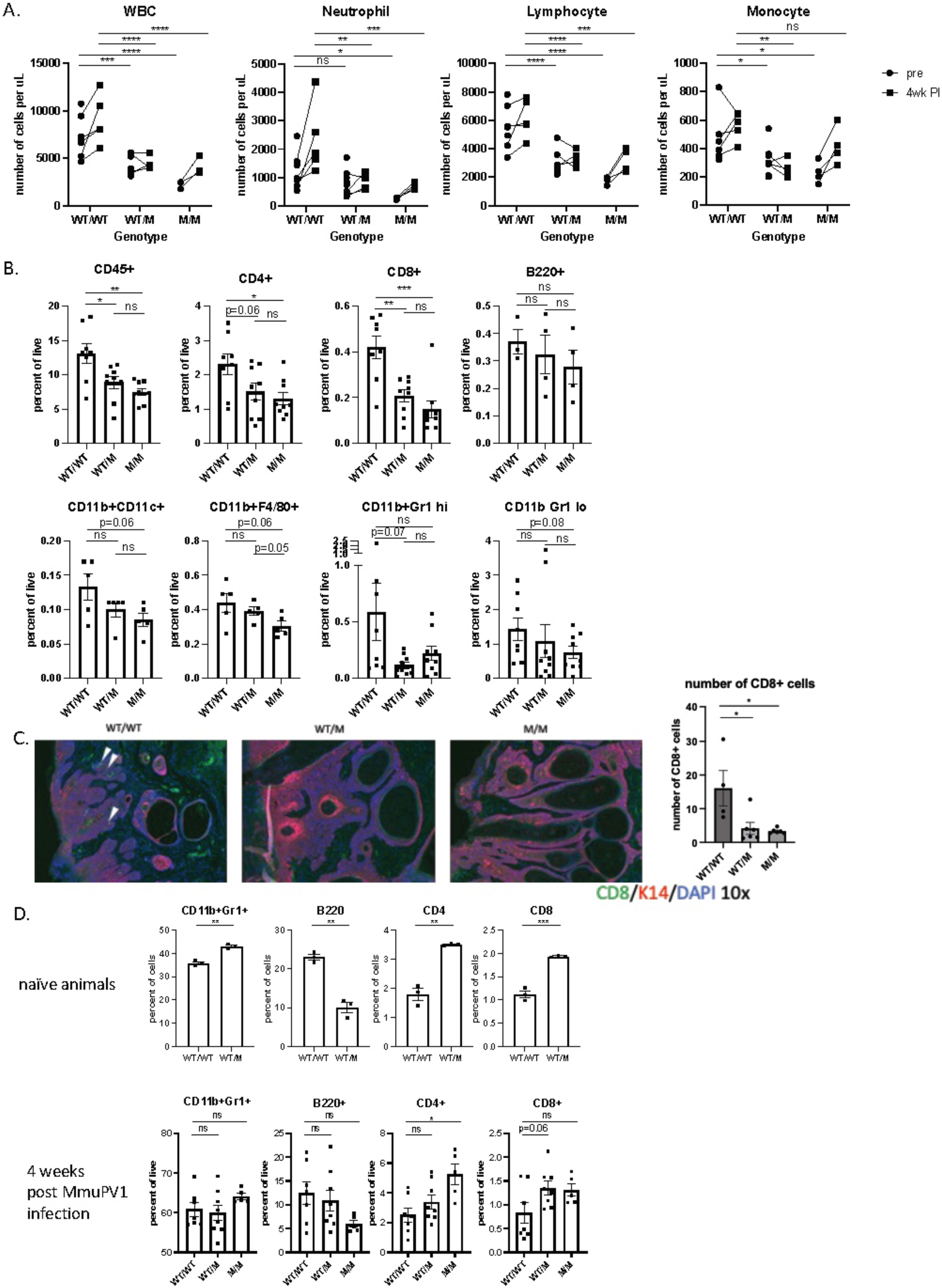
WHIM mice have disturbed T cell distribution in the blood, skin-warts and bone marrow upon MmuPV1 infection. A) Blood were collected from naïve mice and infected mice 4 weeks post infection. Connected data points were from the same mice pre- and post-infection. Blood samples were subject to automatic blood analyzer for circulating white blood cell count (WBC), neutrophil count, lymphocyte count and monocyte count. Two-way ANOVA mixed effect model was used for statistical analysis comparing WT/WT vs. WT/M and WT/WT vs. M/M groups. B) Warts were collected for flow cytometry analysis at 4 weeks post infection. Student t test was used for comparing Infiltrating immune cells (CD45+), CD4+ cells, CD8+ cells, B cells (B220+), neutrophils (CD11b+Gr high), monocytes (CD11b+Gr1 low), macrophages (CD11b+F4/80+) and dendritic cells (CD11b+CD11c+). Student t test was used. C) Immunofluorescence staining of K14+ epithelial cells (red) and CD8+ cells (green). 4-5 fields of 10x images were taken for each wart. Numbers of green cells within red stained epithelia regions were counted for each field and averaged for each wart. Total of 4-6 warts were quantified per genotype. Student t test was used. D) Bone marrow cells were collected for flow cytometry analysis from naïve animals or at 4 weeks post infection. Student t test was used. ns: not significant, *p<0.05, **p<0.01, ***p<0.005, ****p<0.001. Error bars represent standard error of the mean (SEM).

### WHIM bone marrow reconstitution in wildtype mice increases their susceptibility to MmuPV1-induced papillomatosis

It has been reported that CXCL12 is abnormally expressed in HPV-associated lesions (3), and that in the context of the WHIM, there is a potential synergistic role of the gain of CXCR4 function mutation accounting for the tumorigenesis of high risk HPV-immortalized keratinocytes (34). Because we observed lower immune cell infiltration in warts of WHIM mice and higher susceptibility to MmuPV1-induced papillomatosis and wart growth, we sought to test whether the *CXCR4* mutation in hematopoietic cells could confer vulnerability to MmuPV1-mediated papillomatosis. Wildtype FVB/N mice (CD45.2) were subject to lethal dose irradiation (10 Gy) and received whole bone marrow cells from WHIM donor mice (WT/M, CD45.1) or wild type donor mice (WT/WT, CD45.2). Donor bone marrow cells took about 7 weeks to fully reconstitute the recipient mice (Supplemental Figure 1). Seven weeks post bone marrow transplant, blood from recipient mice were sampled for blood cell count and donor chimerism (Supplemental Figure 2). Ears of recipient mice were then infected with MmuPV1 (10^6^ VGE) and warts were scored and measured 4 weeks after MmuPV1 infection (Figure 3A). The baseline level of circulating immune cells were the same among recipients prior to bone marrow transplant (Figure 3B). At 7 weeks post bone marrow transplant, circulating white blood cells including neutrophils, monocytes, CD8+ T cells and B cells were lower in recipients that received bone marrow cells from WHIM donors prior to infection, and stayed low 4 weeks post infection (Figure 3C). In contrast, recipients reconstituted with wildtype bone marrow cells had elevated white blood cell counts upon MmuPV1 infection (4 weeks post infection versus pre-infection) (Figure 3C). Correlated with lower number of circulating white blood cells was the high incidence of papillomatosis (87%) in recipients that received WHIM (WT/M) bone marrow cells. This incidence is significantly higher than the incidence (40%) in mice that received wild type (WT/WT) bone marrow cells (Figure 3D). The average size of warts was also bigger in WHIM bone marrow reconstituted mice (Figure 3E). Half of the warts that developed in recipients with WT bone marrow regressed between week 4 to week 12 post infection (2 out of 4 sites), versus 22% in recipients with WHIM bone marrow (2 out 9 sites) (Figure 3F). Due to low number of warts in recipients with WT bone marrows to begin with, this difference was not significant by Fisher’s exact test. Flow analysis at 4 weeks after infection showed lower infiltrating CD8+ T cells and monocytes from warts arising in WHIM bone marrow reconstituted mice versus wild type bone marrow reconstituted mice (Figure 3G). Our results suggest that WHIM mutation in hematopoietic cells can drive an increase in susceptibility to papillomavirus pathogenesis, correlated with lower circulating white blood cells and lower CD8+ T cell infiltration in warts. The papilloma incidence in WT animals that received radiation and reconstituted with WT bone marrow (Figure 3D) is higher than the incidence in unirradiated WT mice (Figure 1A, 10^6^ VGE). This increased papillomatosis observed in irradiated animals may be a consequence of radiation-induced inflammation (35, 36).

**Figure 3.**
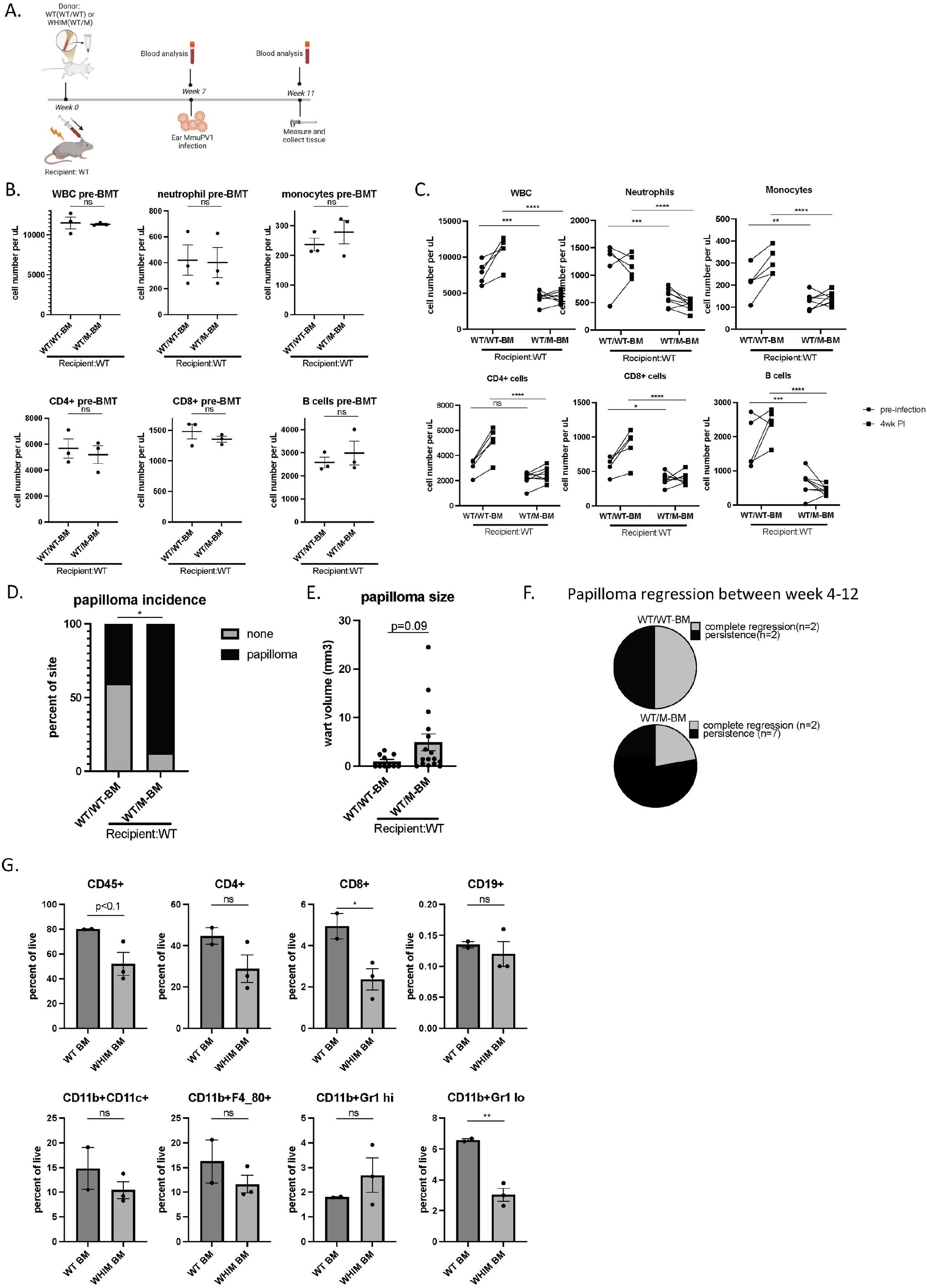
WHIM bone marrow reconstitution in wildtype mice increases their susceptibility to MmuPV1-induced papillomatosis. A) A schema showing timeline for bone marrow transplant and infection. B) Blood were collected prior to bone marrow transplant for white blood cell count (WBC) and subset of immune cells were analyzed by flow cytometry based on percentage of total CD45+ cells times white blood cell count. Student t test was used. C) Blood were collected prior to infection (i.e., 7 weeks post BMT) and 4 weeks post infection for blood count and flow cytometry analysis. Two-way ANOVA was used. D) Papillomatosis incidence was score at 4 weeks post infection. Fisher’s exact test was used. E) Wart size were measured at 4 weeks post infection. Student t test was used. F) Part of the developed papillomas at 4 weeks were followed up until 12 weeks post infection, n=4 papilloma sites from recipients with WT bone marrow, n=9 papilloma sites from recipients with WHIM bone marrow. Pie charts showed number of complete regressed warts versus persisting warts in two groups of recipients. G) Warts were collected at 4 weeks post infection for infiltrating immune cell analysis by flow cytometry. Student t test was used for comparing Infiltrating immune cells (CD45+), CD4+ cells, CD8+ cells, B cells (CD19+), neutrophils (CD11b+Gr high), monocytes (CD11b+Gr1 low), macrophages (CD11b+F4/80+) and dendritic cells (CD11b+CD11c+). ns: not significant, *p<0.05, **p<0.01, ***p<0.005, ****p<0.001. Error bars represent standard error of the mean (SEM).

### Bone marrow transplant from wildtype donors decreases susceptibility to MmuPV1-induced papillomatosis in WHIM mice

Because the WHIM mutation in hematopoietic cells drives susceptibility to MmuPV1 pathogenesis, we next tested whether bone marrow transplant of wildtype bone marrow cells into WHIM mice rescues their phenotype. WHIM mice (WT/M, CD45.1) or wildtype mice (WT/WT, CD45.1) underwent 10 Gy lethal irradiation and then received whole bone marrow cells from wildtype mice (WT/WT, CD45.2) (Figure 4A). Donor chimerism in blood was confirmed 7 weeks after bone marrow transplant, prior to infection (Supplemental Figure 3). Prior to bone marrow transplant, the WHIM recipient mice (WT/M) were confirmed to have lower circulating white blood cells (Figure 4B). At 7 weeks post bone marrow transplant from wild type donors, WHIM recipient mice had similar numbers of circulating white blood cells, including neutrophils, monocytes, CD4 and CD8 T cells, and B cells at both pre-infection and 4 weeks post infection time points compared to wild type recipients (Figure 4C). The circulating neutrophil count was generally high in uninfected animals after radiation (Supplemental Figure 1C) suggesting there was systemic inflammation induced by radiation (35, 36). We did not observe a further increase of circulating neutrophils upon infection in irradiated animals. The incidence of papillomatosis and size of warts were also comparable between WHIM and wild type recipients after bone marrow reconstitution (Figure 4D and 4E). The numbers of wart infiltrating immune cells were also similar between WHIM and wildtype recipients, with the exception for neutrophils (Figure 4F). Despite similar circulating neutrophil numbers, the number of neutrophils infiltrating into warts were lower in WHIM recipients (Figure 4F). However, this difference in level of neutrophil infiltration did not result in any difference in wart size (Figure 4E) or papillomatosis incidence (Figure 4D). Our results indicate that bone marrow transplant is a curative strategy to not only correct neutropenia and lymphopenia in WHIM mice, but also lower their susceptibility to papillomavirus pathogenesis.

**Figure 4.**
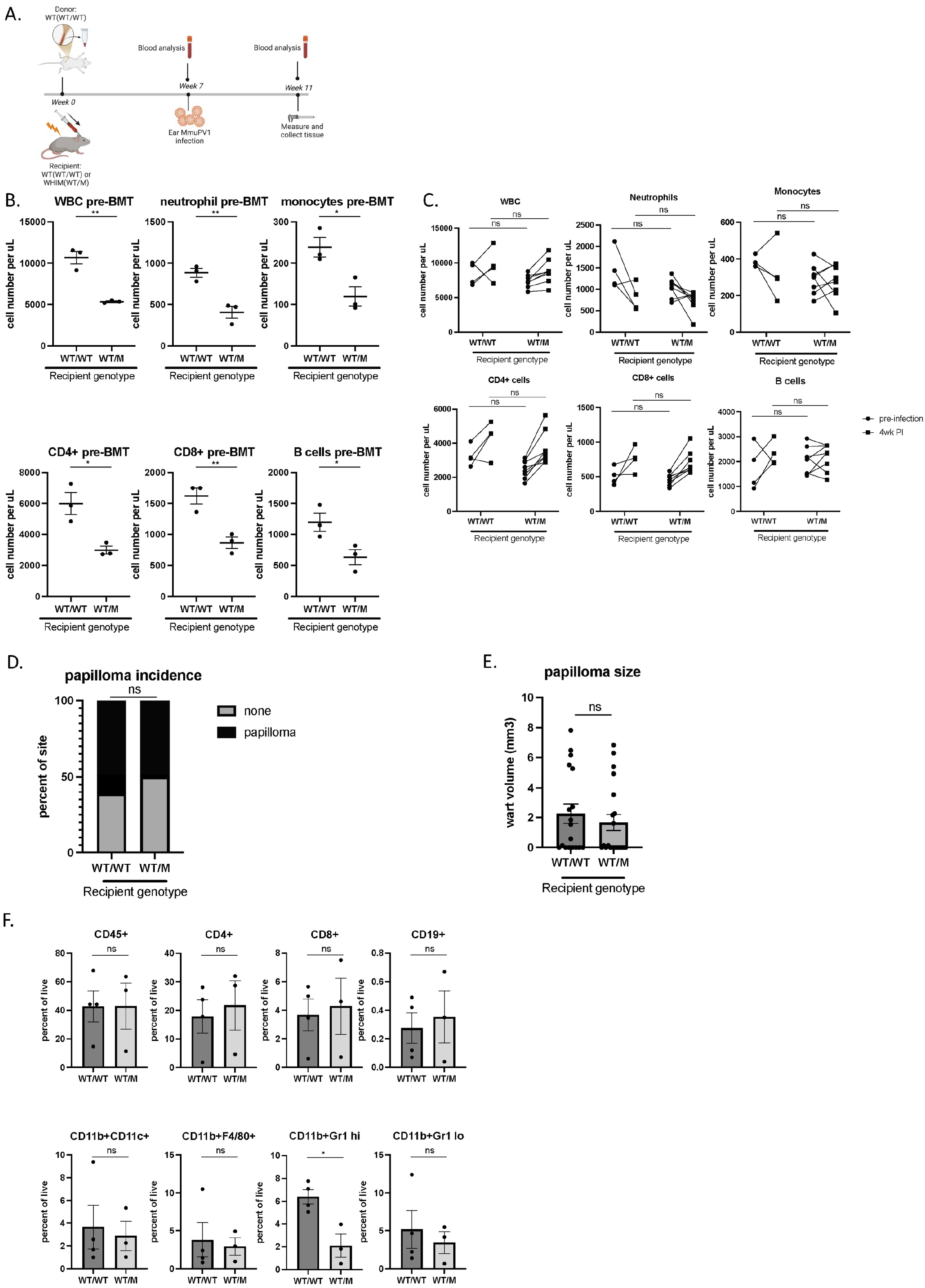
Bone marrow transplant from wildtype donors recues susceptibility of WHIM mice to MmuPV1 induced papillomatosis. A) A schema showing timeline for bone marrow transplant and infection. B) Blood were collected prior to bone marrow transplant for white blood cell count (WBC) and subset of immune cells were analyzed by flow cytometry based on percentage of total CD45+ cells times white blood cell count. Student t test was used. C) Blood was collected prior to infection (i.e. 7 weeks post-BMT) and 4 weeks post infection for blood count and flow cytometry analysis. Two-way ANOVA was used. D) Papillomatosis incidence was score at 4 weeks post infection. Fisher’s exact test was used. E) Wart size were measured at 4 weeks post infection. Student t test was used. F) Warts were collected at 4 weeks post infection for infiltrating immune cell analysis by flow cytometry. Student t test was used for comparing Infiltrating immune cells (CD45+), CD4+ cells, CD8+ cells, B cells (CD19+), neutrophils (CD11b+Gr high), monocytes (CD11b+Gr1 low), macrophages (CD11b+F4/80+) and dendritic cells (CD11b+CD11c+). ns: not significant, *p<0.05, **p<0.01. Error bars represent standard error of the mean (SEM).

## DISCUSSION

Current standard treatments of patients with WHIM syndrome aim to alleviate the symptoms, including granulocyte colony-stimulating factor (G-CSF) or granulocyte–macrophage colony-stimulating factor (GM-CSF) injections to improve neutrophil counts, and monthly infusion of immunoglobulins (7). CXCR4 antagonist, which holds great promise in clinical trials, requires daily administration (13). Therefore, none of these treatments works as a cure for WHIM patients without prolonged treatment (8, 15). A few WHIM patients who underwent allogenic hematopoietic stem cell transplant (HSCT) appeared to be symptom free (14). So far, allogeneic HSCT remains the only curative treatment for WHIM patients, however, it is not integrated into standardized therapy due to high post-HSCT mortality rate and complications. Interestingly, one patient was cured by spontaneous deletion of the WHIM allele (as well as 163 other genes) by chromothripsis in a single hematopoietic stem cell, which repopulated the patient’s whole myeloid repertoire (37). This patient had spontaneous regression of warts and no recurrent infections. This case and our results described in this report provide evidence that WHIM mutations in hematopoietic cells are largely driving the vulnerability to papillomavirus pathogenesis in WHIM patients.

A cell-autonomous effect of the WHIM mutation on keratinocyte transformation has been reported previously (34, 38). Our data showed that warts from WHIM mice had a significantly higher (3-4 fold) number of cells bearing amplified MmuPV1 DNA 4 weeks post infection (Figure 1D), though this did not correlate with higher levels of viral RNA or proteins (Figures 1E and 1F). This finding indicates that keratinocytes carrying the WHIM mutation are more capable of supporting the productive phase of viral DNA replication, because the strong, nuclear, *in situ* hybridization signal is thought to be reflective of the viral DNA amplification that is associated with the productive phase of the viral life cycle. This difference may reflect a cell-autonomous effect of the WHIM mutation on keratinocyte biology and/or a cell non-autonomous effect of the WHIM mutation on the microenvironment within the wart. Further studies are required to study the underlying mechanisms.

The mouse strain bearing human WHIM mutation allele has been useful to test the potential of therapeutic drugs and gene therapy to reverse leukopenia in WHIM syndrome (31, 32, 39). However, whether the reversion of leukopenia is enough to prevent future HPV pathogenesis and which immune cells are important to control papillomavirus infections remain unclear. Using murine papillomavirus (MmuPV1) as an infection model in mice, we demonstrate that the WHIM mutation *in hematopoietic cells* increased susceptibility to papillomavirus pathogenesis and that correction of leukopenia by bone marrow transplant correlated with prevention of new papillomatosis. We also found a potential cell-autonomous effect of WHIM on increasing the amplification of viral DNA within keratinocytes of warts.

The single patient with a spontaneous cure by chromothripsis had restored myeloid cell counts but retained lymphopenia during and after her warts resolved (37). This suggests that myeloid cells play a critical role in facilitating wart regression. Here we show that local MmuPV1 infection increases circulating white blood cell count including both myeloid and lymphoid cells in wild type animals but not in WHIM animals (Figure 2A). A transient increase may have occurred earlier after infection in WHIM animals, but it is hard to distinguish such a response from a response of wound healing process that takes place in the first three weeks after scarring/infection. However, in the warts developed in WHIM mice, we observed no significant drop of infiltrating myeloid cells as compared to wild type animals (Figure 2B) despite a significant difference of myeloid cells in circulation (Figure 2A). In contrast, wart-infiltrating CD4 and CD8 T cells were significantly diminished in WHIM mice (Figure 2B and 2C). In mice reconstituted with WHIM bone marrow cells, again we observed lower wart-infiltrating CD8 T cells (Figure 3G), correlating with increased wart growth and incidence (Figure 3D and 3E). Our data indicate that T cells are likely critical for papillomatosis control upon MmuPV1 infection, as reported previously (24, 25, 40, 41). This finding is also consistent with other reports that in HPV-associated human cancers, T cell response is associated with better clinical outcomes (42–45), and HPV-prevalence of types associated with mucosal lesion is higher in HIV-positive patients who have deficiency in T cell response (46, 47). Functional HPV-specific stem-like CD8 T cells are present in HPV-positive head and neck cancer, supporting an important role of CD8 T cells in HPV associated diseases (48).

Our results demonstrate that the WHIM mouse strain is a powerful pre-clinical model to develop infectious and malignant disease models for understanding HPV-host interactions and testing treatment options for WHIM syndrome. Recent efforts investigating gene therapy as a curative treatment for WHIM syndrome have shown that leukopenia correction can be achieved in unconditioned WHIM mice by CXCR4-haploinsufficient bone marrow transplant (39). Although it is attractive to reconstitute WHIM recipients without prior conditioning, the reconstitution of donor T cells in circulation is poor compared to myeloid cells and B cells (39). Future investigations are needed to test whether it is possible to prevent papillomavirus pathogenesis by reconstituting unconditioned WHIM recipients. In parallel, long-term follow up should be warranted to test if transplant of wild type or genetically corrected HSCs can cure established malignancies caused by papillomavirus.

## MATERIALS AND METHODS

### Animals

WHIM mice on FVB/N background were provided by Dr. Francoise Bachelerie (French Institute of Health and Medical Research) (31). Wildtype FVB/N mice were obtained from Taconic and used as breeders to generate experimental mice. WHIM mice that were heterozygous for WHIM mutation (CXCR4^1013/+^) were mated to Wildtype FVB/N or WHIM mice (CXCR4^1013/+^) to generate WHIM mice carrying one (WT/M) or two (M/M) CXCR4 mutant genes. Wildtype littermates were used as WT controls for each experiment. All mice were housed in animal facility in aseptic conditions in micro-isolator cages and experiments carried out under an approved animal protocol. Six-to ten-week-old mice were used for experiments with a same ratio of male and females in each group.

### Ethics statement

Experiments were approved and performed in accordance with guidelines approved by the Association for Assessment of Laboratory Animal Care, at the University of Wisconsin Medical School. This study was approved by the University of Wisconsin School of Medicine and Public Health Institutional Animal Care and Use Committee under protocol number M005871.

### Bone marrow transplant

Recipient mice were subject to lethal dose whole body irradiation (10 Gy). Recipients then received 8 million whole bone marrow cells from congenic donor mice on the same day of radiation. Submandibular bleeding was performed at indicated time points to confirm chimerism in recipient mice. Seven weeks post bone marrow transplant, mice were infected on ears with MmuPV1.

### MmuPV1 isolation and infection

Ears of nude mice were scarified and infected with prior MmuPV1 stock virus at 1x10^8^ VGE/ul. Warts from ears were collected 3 months after infection for crude viral preparations. In brief, warts were homogenized in PBS and incubated at 37C for 30 min in the presence of benzonase and triton. Wart suspensions were then digested with collagenase IV at 4°C for 48 hours. Suspensions were spun down and supernatant were collected and aliquoted as rude virus stock. 10ul of Virus stock was digested with proteinase K and loaded onto agarose gel in multiple dilutions to quantify viral genome equivalent (VGE). A plasmid that was of the same size as MmuPV1 was used as standard. For experimental infection using prepared virus stock, virus stock with a determined VGE was diluted to various concentrations described for each experiment. Mouse ears were scarified and wounded using a 27-gauge needle before infection. 2 ul of virus stock with indicated VGE in each figure legend were applied to the scarified ears. Papilloma width, length and height were measured by caliper and used for calculation of papilloma size (width x length x height).

### Complete blood cell count

50-100 ul of blood was collected for each mouse by submandibular bleeding in EDTA coated tubes. Complete blood counts, neutrophil and lymphocyte counts were determined by Hemavet auto blood analyzer. 10-20 ul of blood per mouse was processed for flow cytometry analysis if subpopulation of white blood cell analysis was performed.

### Tissue collection for histology analysis

Warts were cut in half and fixed in 4% PFA at 4°C overnight and switched to 70% ethanol. They were then embedded in paraffin and sectioned into 5 microns pieces. For H&E staining, tissue sections were deparaffinized and rehydrated by going through gradient ethanol solutions and stained with hematoxylin and eosin. Tissues sections were then dehydrated and mounted in Cytoseal media.

### Immunofluorescent staining

Tissues were deparaffinized in Xylene twice and rehydrated in gradient ethanol. Antigen retrieval was performed in boiling citrate buffer (pH=6) for 20 min. Tissues were blocked in 3% hydrogen peroxide followed by TSA blocking buffer (Perkin Elmer #FP1012). Tissues were then incubated with rabbit Sera against MmuPV1 L1 (provided by Dr. Chris Buck, National Cancer Insitute, Bethesda, MD) overnight at 4°. Goat anti-rabbit HRP antibody (Invitrogen #31420) was applied on tissue for 1 hour at room temperature, followed by tyramide-biotin complex amplification for 10 min at room temperature. Rabbit Sera against MmuPV1 E4 (provided by Professor John Doorbar, University of Cambridge, UK) was applied on tissue for overnight at 4°. Goat anti-rabbit Alexa488 (Molecular Probes A11108) and Alexa647 conjugated streptavidin were applied for 1 hour at room temperature. Tissues were then counterstained with Hoechst dye and mounted and cover slipped.

### *In situ* hybridization

MmuPV1 viral DNA and RNA were detected using RNAscope 2.5 HD Assay-Brown (Advanced Cell Diagnostics, Newark, CA) according to the manufacturer’s instructions with probes specific for MmuPV1 E1∧E4 (catalog no. 473281). For DNA detection, tissues were treated with RNase A and RNase I (Thermos Fisher). For RNA detection, tissues were treated with RNase-free DNase (Thermo Fisher). Slides were counterstained with hematoxylin before mounting and cover slipping.

### Quantification of MmuPV1 DNA, RNA and proteins

20X images were taken for 3 fields per mouse wart. The number of cells positive for E4 protein and L1 protein were counted for each field and averaged for each mouse wart. Mean fluorescence of E4 protein and L1 protein were measured in ImageJ for each field and averaged for each mouse wart. DNA or RNA positive cells and total nuclei numbers were quantified by an automatic counting program “BrdU Count v2.1.ijm” developed by Dr. David Ornelles (Wake Forest University).

### Flow cytometry

Mouse whole blood and bone marrow samples was lysed using red blood cell lysis buffer (Tonbo biosciences) for 10 min at room temperature, washed in PBS and then stained with conjugated antibodies for immune cell surface markers at 4° C for 30 min. Cell suspensions were washed in PBS with 5% FBS and analyzed by ThermoFisher Attune. Ear warts were trimmed of surrounding tissues and harvested on ice in PBS. Warts were cut into 1 mm pieces and digested in RPMI supplemented with Miltenyie mouse tumor dissociation kit at 37°C for 30 min. Tissues were then homogenized with the back of 1 ml syringe, passed through 70 μM filter and washed twice with cold PBS. Single cell suspensions were then stained with 1 ul Zombie live/dead staining dye (Biolegend) in 500ul pf PBS at 4°C for 15 min. Cells were then washed with PBS supplemented with 5% FBS, and stained with cell surface markers. Wart cell suspensions were then washed and fixed with fixation buffer (eBioscience) overnight in 4°C. Cells were washed in PBS supplemented with 5% FBS and analyzed with ThermoFisher Attune. Compensation beads (eBioscience) stained with each antibody were used as single-color controls. A combination of selected antibodies (anti-mouse) was used depending on the purpose of each study: CD45 APC-Cy7 (Biolegend, clone 30-F11), CD8a FITC (Tonbo ebioscience, clone 53–6.7), CD4 PE (Tonbo ebioscience, clone RM4-5), Gr1 PE-Cy5 (Biole-gend, clone RB6-8C5), F4/80 BV421 (Biolegend clone BM8), CD11b BV605 (Biolegend, clone M1/70), CD11c PE-Cy7 (Biolegend, clone N418).

### Statistics

All statistical analyses were done with Graphpad Prism. Two-way ANOVA was used for statistical comparison when two variables (infection and genotype) were involved. For single variable experiments, t test or one-way ANOVA was used for statistical comparison as indicated.

## ACKNOWLEDGMENTS

The authors thank the University of Wisconsin Translational Research Initiatives in Pathology (TRIP) laboratory, supported by the UW Department of Pathology and Laboratory Medicine, UWCCC (P30 CA014520) and the Office of The Director-NIH (S10OD023526) for use of its facilities and services. We thank Amanda Loke for help in processing samples. We thank Dr. Chris Buck, (National Cancer Insitute, Bethesda, MD) and Professor John Doorbar (University of Cambridge, UK) for providing antibodies. We thank Jing Zhang for her guidance on radiation treatment for mouse bone marrow transplant studies.

This research was supported by the 2020 AACR-Genentech Immuno-oncology Research Fellowship, Grant Number 20–40-18-WANG to W.W., the University of Wisconsin-Madison Carbone Cancer Center Transdisciplinary Cancer Immunology-Immunotherapy Pilot Grant to P.F.L., grants from the National Institutes of Health (P01 CA022443, R35 CA210807, P50 CA278595, R01 CA228543 to P.F.L.) and “Fondation Pour la Recherche Médicale” project N° EQU202203014751 to F.B.

## Supporting information captions

**Supplemental Figure 1.**
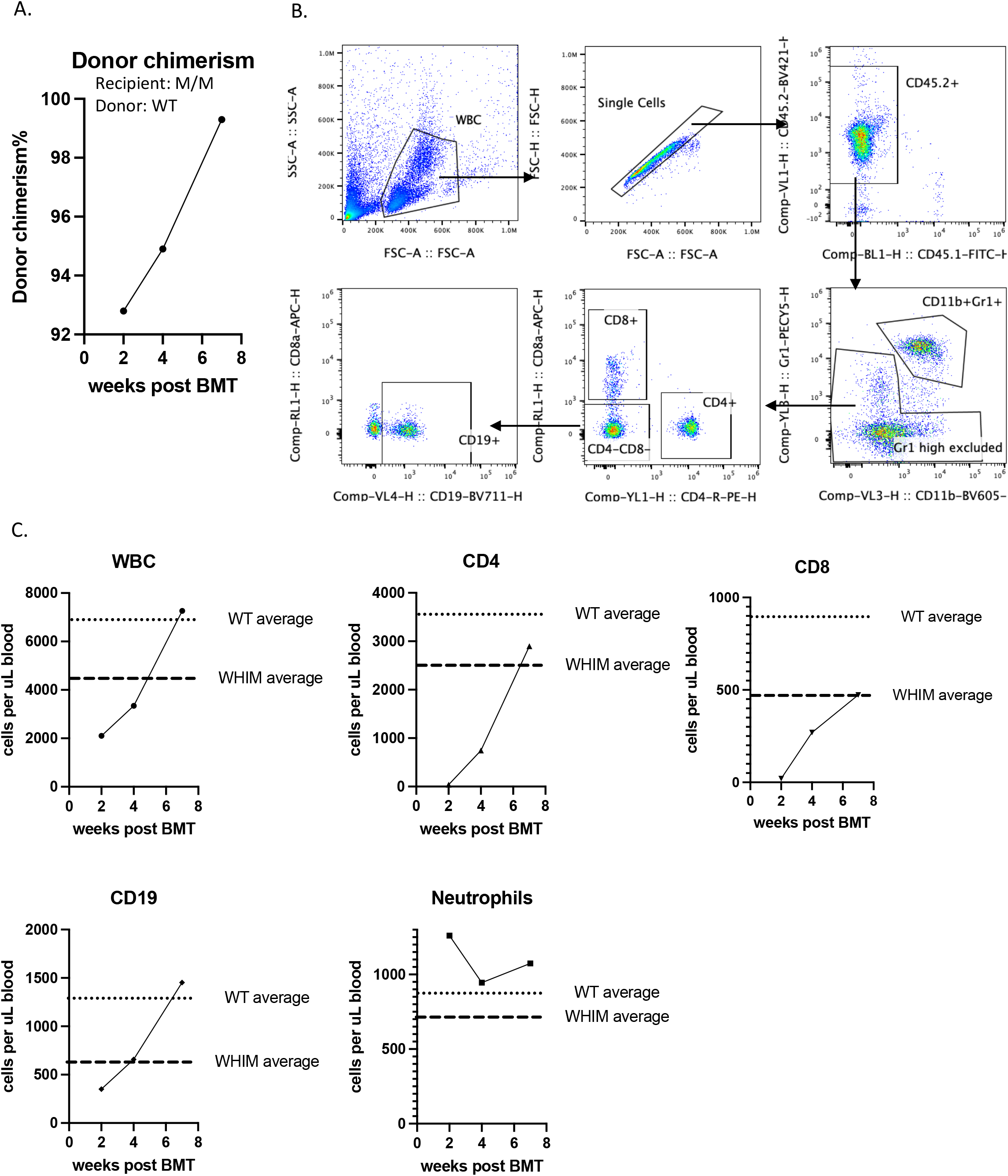
Timeline establishment of donor bone marrow reconstitution in FVB/N mice. WHIM mouse (M/M, CD45.1) received lethal dose 10Gy of total body irradiation. On the same day, the recipient received 8 million total bone marrow cells from a congenic wildtype donor (WT/WT, CD45.2). Blood was collected at 2 weeks, 4 weeks and 7 weeks post bone marrow transplant and were subject to flow cytometry analysis and complete blood count. A) Donor chimerism in circulating blood. B) Gating strategy for flow cytometry analysis. C) White blood cell count (WBC) and subtypes of immune cells in circulating blood of recipient mouse. Dotted lines represent average circulating count from naïve wildtype (WT) animals and WHIM heterozygous (WHIM) animals.

**Supplemental Figure 2.**
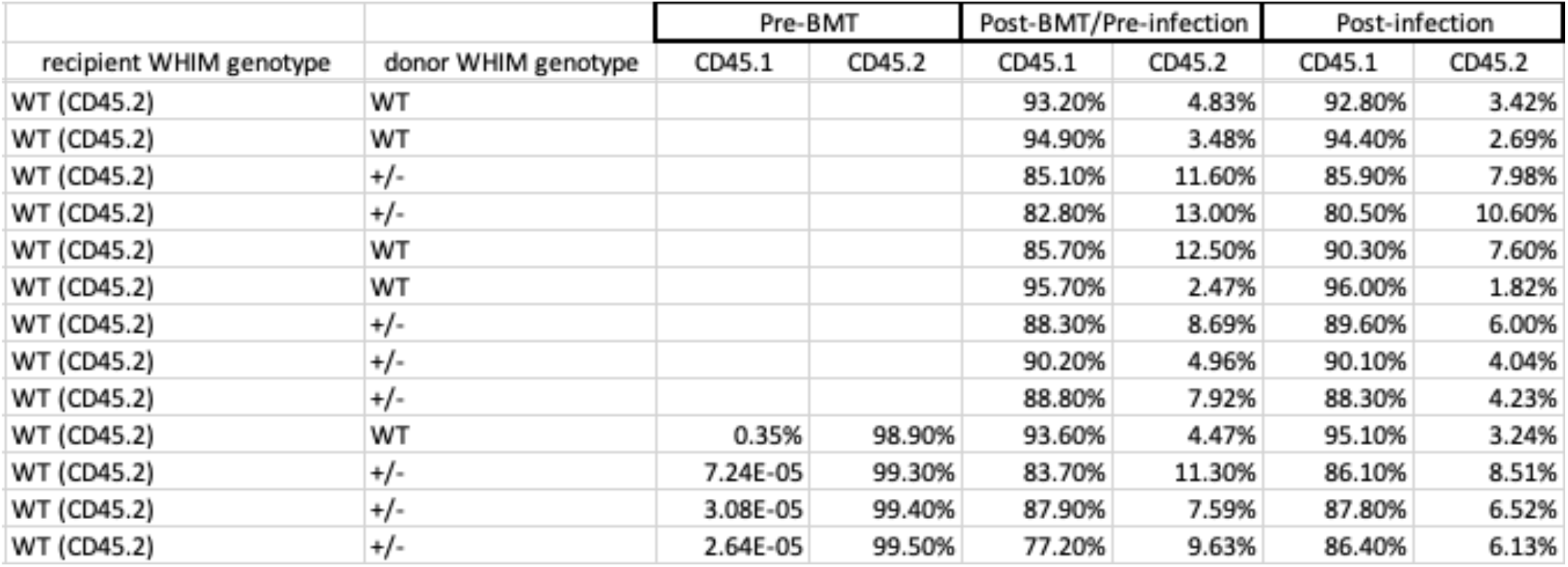
Donor chimerism in recipients. Wildtype recipient mice (WT, CD45.2) received lethal dose 10Gy of total body irradiation. On the same day, the recipients received 8 million total bone marrow cells from a congenic wildtype donor (WT, CD45.1) or a heterozygous WHIM donor mouse (+/-, CD45.1). Blood was collected from a subset of recipients prior to bone marrow transplant (pre-BMT), from all recipient 7 weeks post bone marrow transplant but prior to infection (Post-BMT/Pre-infection), from all recipients 4 weeks post infection (Post-infection) to check donor reconstitution (percentage of CD45.1+ cells) in circulating blood.

**Supplemental Figure 3.**
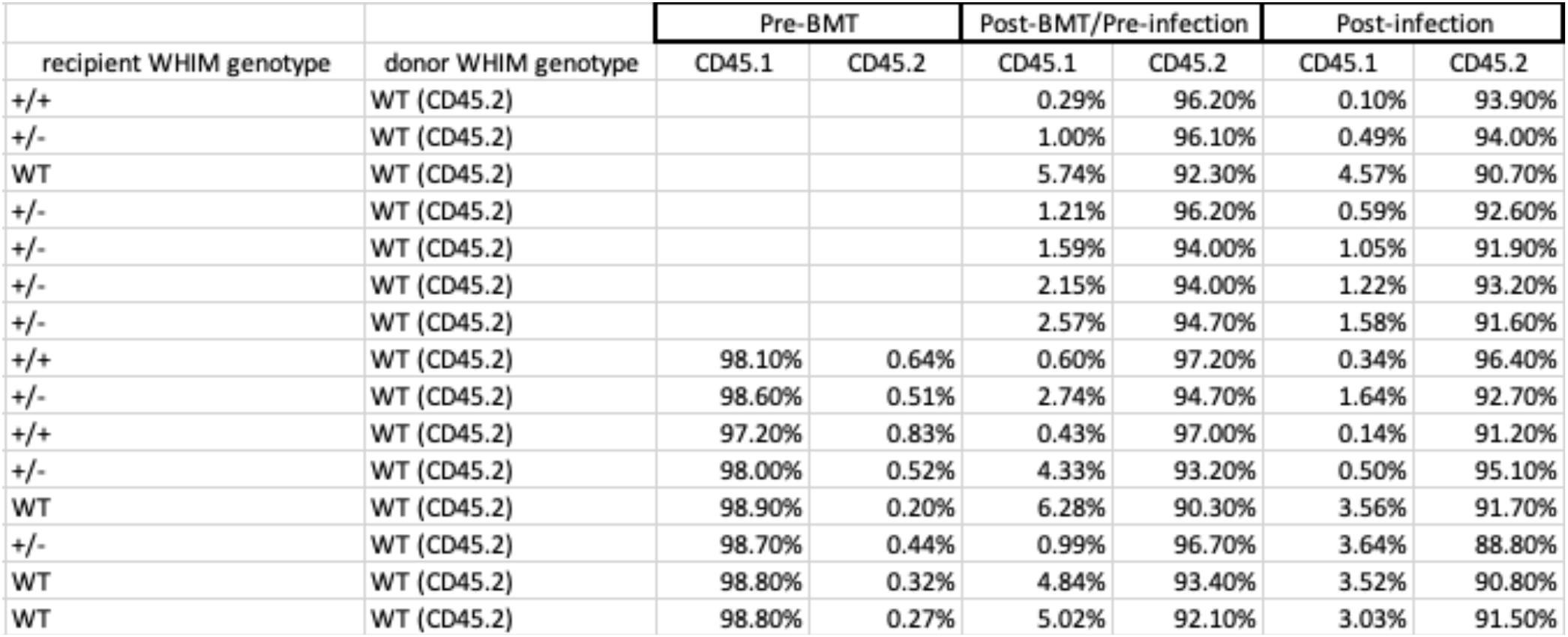
Donor chimerism in recipients. Wildtype mice (WT, CD45.1) or heterozygous WHIM mice (+/-, CD45.1) received lethal dose 10Gy of total body irradiation. On the same day, the recipients received 8 million total bone marrow cells from a congenic wildtype donor (WT, CD45.2). Blood was collected from a subset of recipients prior to bone marrow transplant (pre-BMT), from all recipient 7 weeks post bone marrow transplant but prior to infection (Post-BMT/Pre-infection), from all recipients 4 weeks post infection (Post-infection) to check donor reconstitution (percentage of CD45.2+ cells) in circulating blood.

## REFERENCES

1. Kawai T, Malech HL. WHIM syndrome: congenital immune deficiency disease. Curr Opin Hematol. 2009;16(1):20–6.

2. Hernandez PA, Gorlin RJ, Lukens JN, Taniuchi S, Bohinjec J, Francois F, et al. Mutations in the chemokine receptor gene CXCR4 are associated with WHIM syndrome, a combined immunodeficiency disease. Nat Genet. 2003;34(1):70–4.

3. Balabanian K, Lagane B, Pablos JL, Laurent L, Planchenault T, Verola O, et al. WHIM syndromes with different genetic anomalies are accounted for by impaired CXCR4 desensitization to CXCL12. Blood. 2005;105(6):2449–57.

4. Sugiyama T, Kohara H, Noda M, Nagasawa T. Maintenance of the hematopoietic stem cell pool by CXCL12-CXCR4 chemokine signaling in bone marrow stromal cell niches. Immunity. 2006;25(6):977–88.

5. Nie Y, Han YC, Zou YR. CXCR4 is required for the quiescence of primitive hematopoietic cells. J Exp Med. 2008;205(4):777–83.

6. Susek KH, Karvouni M, Alici E, Lundqvist A. The Role of CXC Chemokine Receptors 1-4 on Immune Cells in the Tumor Microenvironment. Front Immunol. 2018;9:2159.

7. McDermott DH, Murphy PM. WHIM syndrome: Immunopathogenesis, treatment and cure strategies. Immunol Rev. 2019;287(1):91–102.

8. David C. Dale FCF, Audrey Anna Bolyard, Weihua Tang, Honghua Jiang, Richard MacLeod, Diego Cadavid, Yanping Hu. Mavorixafor, an Oral CXCR4 Antagonist, for Treatment of Patients with WHIM Syndrome: Results from the Long-Term Extension of the Open-Label Phase 2 Study. Blood. 2021;138:1121.

9. Bilgin YM. Use of Plerixafor for Stem Cell Mobilization in the Setting of Autologous and Allogeneic Stem Cell Transplantations: An Update. J Blood Med. 2021;12:403–12.

10. Devine SM, Vij R, Rettig M, Todt L, McGlauchlen K, Fisher N, et al. Rapid mobilization of functional donor hematopoietic cells without G-CSF using AMD3100, an antagonist of the CXCR4/SDF-1 interaction. Blood. 2008;112(4):990–8.

11. Meuris F, Gaudin F, Aknin ML, Hemon P, Berrebi D, Bachelerie F. Symptomatic Improvement in Human Papillomavirus-Induced Epithelial Neoplasia by Specific Targeting of the CXCR4 Chemokine Receptor. J Invest Dermatol. 2016;136(2):473–80.

12. McDermott DH, Liu Q, Ulrick J, Kwatemaa N, Anaya-O’Brien S, Penzak SR, et al. The CXCR4 antagonist plerixafor corrects panleukopenia in patients with WHIM syndrome. Blood. 2011;118(18):4957–62.

13. McDermott DH, Pastrana DV, Calvo KR, Pittaluga S, Velez D, Cho E, et al. Plerixafor for the Treatment of WHIM Syndrome. N Engl J Med. 2019;380(2):163–70.

14. Laberko A, Deordieva E, Krivan G, Goda V, Bhar S, Kawahara Y, et al. Multicenter Experience of Hematopoietic Stem Cell Transplantation in WHIM Syndrome. J Clin Immunol. 2022;42(1):171–82.

15. Dotta L, Notarangelo LD, Moratto D, Kumar R, Porta F, Soresina A, et al. Long-Term Outcome of WHIM Syndrome in 18 Patients: High Risk of Lung Disease and HPV-Related Malignancies. J Allergy Clin Immunol Pract. 2019;7(5):1568–77.

16. Beaussant Cohen S, Fenneteau O, Plouvier E, Rohrlich PS, Daltroff G, Plantier I, et al. Description and outcome of a cohort of 8 patients with WHIM syndrome from the French Severe Chronic Neutropenia Registry. Orphanet J Rare Dis. 2012;7:71.

17. Badolato R, Donadieu J, Group WR. How I treat warts, hypogammaglobulinemia, infections, and myelokathexis syndrome. Blood. 2017;130(23):2491–8.

18. Handisurya A, Schellenbacher C, Reininger B, Koszik F, Vyhnanek P, Heitger A, et al. A quadrivalent HPV vaccine induces humoral and cellular immune responses in WHIM immunodeficiency syndrome. Vaccine. 2010;28(30):4837–41.

19. Palm MD, Tyring SK, Rady PL, Tharp MD. Human papillomavirus typing of verrucae in a patient with WHIM syndrome. Arch Dermatol. 2010;146(8):931–2.

20. Pastrana DV, Peretti A, Welch NL, Borgogna C, Olivero C, Badolato R, et al. Metagenomic Discovery of 83 New Human Papillomavirus Types in Patients with Immunodeficiency. mSphere. 2018;3(6).

21. Ingle A, Ghim S., Joh J., Chepkoech I., Jenson A.B.,, P. SJ. Novel Laboratory Mouse Papillomavirus (MusPV) Infection. Veterinary Pathology. 2011;48(2):500–5.

22. Bilger A, King RE, Schroeder JP, Piette JT, Hinshaw LA, Kurth AD, et al. A Mouse Model of Oropharyngeal Papillomavirus-Induced Neoplasia Using Novel Tools for Infection and Nasal Anesthesia. Viruses. 2020;12(4).

23. Blaine-Sauer S, Shin MK, Matkowskyj KA, Ward-Shaw E, Lambert PF. A Novel Model for Papillomavirus-Mediated Anal Disease and Cancer Using the Mouse Papillomavirus. mBio. 2021;12(4):e0161121.

24. Wang W, Uberoi A., Spurgeon M., Gronski E., Majerciak V., Lobanov A., Hayes M., Loke A., Zheng Z.M., Lambert P.F. Stress keratin 17 enhances papillomavirus infection-induced disease by downregulating T cell recruitment. PLoS Pathog. 2020;16(1).

25. Wang W, Spurgeon ME, Pope A, McGregor S, Ward-Shaw E, Gronski E, et al. Stress keratin 17 and estrogen support viral persistence and modulate the immune environment during cervicovaginal murine papillomavirus infection. Proc Natl Acad Sci U S A. 2023;120(12):e2214225120.

26. Wei T, Buehler D, Ward-Shaw E, Lambert PF. An Infection-Based Murine Model for Papillomavirus-Associated Head and Neck Cancer. mBio. 2020;11(3).

27. DePianto D, Kerns M., Dlugosz A.A., Coulombe P.A. Keratin 17 promotes epithelial proliferation and tumor growth by polarizing the immune response in skin. Nature Genetics. 2010;42:910–4.

28. Hobbs RP, Batazzi A.S., Han M.C., Coulombe P.A. Loss of Keratin 17 induces tissue-specific cytokine polarization and cellular differentiation in HPV16-driven cervical tumorigenesis in vivo. Oncogene. 2016;35(43):5653–62.

29. Zhussupbekova S, Sinha R., Kuo P., Lambert P.F., Frazer I.H., Tuong Z.K. A Mouse Model of Hyperproliferative Human Epithelium Validated by Keratin Profiling Shows an Aberrant Cytoskeletal Response to Injury. EBioMedicine. 2016;9: 314–23.

30. Wang W, Lozar T, Golfinos AE, Lee D, Gronski E, Ward-Shaw E, et al. Stress Keratin 17 Expression in Head and Neck Cancer Contributes to Immune Evasion and Resistance to Immune-Checkpoint Blockade. Clin Cancer Res. 2022;28(13):2953–68.

31. Balabanian K, Brotin E, Biajoux V, Bouchet-Delbos L, Lainey E, Fenneteau O, et al. Proper desensitization of CXCR4 is required for lymphocyte development and peripheral compartmentalization in mice. Blood. 2012;119(24):5722–30.

32. Gao JL, Owusu-Ansah A, Yang A, Yim E, McDermott DH, Jacobs P, et al. CRISPR/Cas9-mediated Cxcr4 Disease Allele Inactivation for Gene Therapy in a Mouse Model of WHIM Syndrome. Blood. 2023.

33. Uberoi A, Yoshida S, Lambert PF. Development of an in vivo infection model to study Mouse papillomavirus-1 (MmuPV1). J Virol Methods. 2018;253:11–7.

34. Chow KY, Brotin E, Ben Khalifa Y, Carthagena L, Teissier S, Danckaert A, et al. A pivotal role for CXCL12 signaling in HPV-mediated transformation of keratinocytes: clues to understanding HPV-pathogenesis in WHIM syndrome. Cell Host Microbe. 2010;8(6):523–33.

35. Schaue D, Micewicz ED, Ratikan JA, Xie MW, Cheng G, McBride WH. Radiation and inflammation. Semin Radiat Oncol. 2015;25(1):4–10.

36. Stoecklein VM, Osuka A, Ishikawa S, Lederer MR, Wanke-Jellinek L, Lederer JA. Radiation exposure induces inflammasome pathway activation in immune cells. J Immunol. 2015;194(3):1178–89.

37. McDermott DH, Gao JL, Liu Q, Siwicki M, Martens C, Jacobs P, et al. Chromothriptic cure of WHIM syndrome. Cell. 2015;160(4):686–99.

38. Meuris F, Carthagena L, Jaracz-Ros A, Gaudin F, Cutolo P, Deback C, et al. The CXCL12/CXCR4 Signaling Pathway: A New Susceptibility Factor in Human Papillomavirus Pathogenesis. PLoS Pathog. 2016;12(12):e1006039.

39. Gao JL, Yim E, Siwicki M, Yang A, Liu Q, Azani A, et al. Cxcr4-haploinsufficient bone marrow transplantation corrects leukopenia in an unconditioned WHIM syndrome model. J Clin Invest. 2018;128(8):3312–8.

40. Handisurya A. DPM, Thompson C.D., Bonelli M., Lowy D.R., Schiller J.T. Strain-specific properties and T cells regulate the susceptibility to papilloma induction by Mus musculus papillomavirus 1. PLoS Pathog. 2014;10(8):e1004314.

41. Wang J.W. JR, Peng S., Chang Y.N., Hung C.F., Roden R.B. Immunologic Control of Mus musculus Papillomavirus Type 1. PLoS Pathog. 2015;11(10):e1005243.

42. Bhatt KH, Neller MA, Srihari S, Crooks P, Lekieffre L, Aftab BT, et al. Profiling HPV-16-specific T cell responses reveals broad antigen reactivities in oropharyngeal cancer patients. J Exp Med. 2020;217(10).

43. Cai H, Feng Y, Fan P, Guo Y, Kuerban G, Chang C, et al. HPV16 E6-specific T cell response and HLA-A alleles are related to the prognosis of patients with cervical cancer. Infect Agent Cancer. 2021;16(1):61.

44. Kortekaas KE, Santegoets SJ, Abdulrahman Z, van Ham VJ, van der Tol M, Ehsan I, et al. High numbers of activated helper T cells are associated with better clinical outcome in early stage vulvar cancer, irrespective of HPV or p53 status. J Immunother Cancer. 2019;7(1):236.

45. Mondatore D, Bai F, Augello M, Giovenzana M, Pisani Ceretti A, Bono V, et al. Persistence of High Percentage of Peripheral Activated CD8+ T Cells Predict Cytologic HPV-Related Dysplasia in cART-Treated, HIV-Positive Subjects. Open Forum Infect Dis. 2022;9(4):ofac046.

46. Aziz H, Sattar AA, Mahmood H, Fatima S, Khurshid M, Faheem M. Prevalence of HPV types in HIV-positive and negative females with normal cervical cytology or dysplasia. J Clin Lab Anal. 2023;37(4):e24851.

47. Tartaglia E, Falasca K, Vecchiet J, Sabusco GP, Picciano G, Di Marco R, et al. Prevalence of HPV infection among HIV-positive and HIV-negative women in Central/Eastern Italy: Strategies of prevention. Oncol Lett. 2017;14(6):7629–35.

48. Eberhardt CS, Kissick HT, Patel MR, Cardenas MA, Prokhnevska N, Obeng RC, et al. Functional HPV-specific PD-1(+) stem-like CD8 T cells in head and neck cancer. Nature. 2021;597(7875):279-84.

